# An Experimental Test of Local Adaptation in Native and Introduced Populations of an Ectomycorrhizal Fungus, *Suillus luteus*

**DOI:** 10.1101/2024.09.14.613092

**Authors:** Brooke M. Allen, Rytas J. Vilgalys, Jason D. Hoeksema

## Abstract

After species introductions and subsequent invasions, organisms often encounter intense selection pressures from biotic and abiotic aspects of novel environments, driving rapid evolutionary changes that potentially lead to local adaptation. This study aimed to investigate how invasion has influenced symbiotic interactions through rapid evolution in exotic isolates of the ectomycorrhizal (EcM) fungus *Suillus luteus*, which was co-introduced with obligately symbiotic pine trees into the Southern Hemisphere. We conducted a cross-inoculation experiment testing the compatibility of sympatric and allopatric pairings between pines and isolates of *S. luteus* from native and introduced populations. Our results showed that plant and fungal performance were substantially affected by these pairings, largely supporting a hypothesis of local maladaptation. Several performance metrics indicated stronger outcomes in allopatric pairings compared to sympatric ones. This suggests that fungal isolates may have evolved traits that are less beneficial or even somewhat harmful to their local host plants. These findings highlight the complex dynamics of coevolution and emphasize the necessity of considering both local adaptation and maladaptation in understanding species interactions.

## 1. Introduction

After species introductions and during subsequent invasions, organisms often encounter intense selection pressures from biotic and abiotic aspects of novel environments (Osbourn, 2010; Colautti and Lau, 2016). These pressures can drive rapid evolutionary changes, potentially leading to local adaptation, when natural selection favors genotypes that enhance fitness in the invaded habitat (Thompson et al., 2002; Kraemer and Boynton, 2017). These evolutionary processes unfold swiftly, particularly in organisms with short generation times, significantly impacting ecological dynamics, species interactions, and community function. Despite their crucial role, relatively few studies have quantified the extent of rapid evolution in biological invasions, especially in mutualistic species interactions (Thompson, 2013; Colautti and Lau, 2016). This study aimed to investigate how invasion has influenced symbiotic interactions through rapid evolution in exotic isolates of the ectomycorrhizal (EcM) fungus *Suillus luteus. Suillus luteus* was co-introduced with obligately symbiotic pine trees into the Southern Hemisphere, providing a unique opportunity to explore ecological and evolutionary dynamics in this context.

Mycorrhizae are an example of nutritional mutualism, where fungi form symbiotic associations with plant roots, providing soil nutrients to the plant in exchange for fixed carbon (Smith and Read, 2010). Ectomycorrhizal (EcM) fungi predominantly associate with trees, many of which, like obligately ectomycorrhizal pines (*Pinaceae*), hold ecological or economic significance. Historically, from the mid-1800s to the 1940s, *Pinus contorta, Pinus pinaster*, and *Pinus radiata* were introduced across the Southern Hemisphere from North America and Eurasia to establish pine plantations (Critchfield et al., 1966; Scott, 1960; Australian Bureau of Agricultural and Resource Economics, 2008). Alongside these pines came their EcM fungal symbionts, essential for plantation success (Richardson et al., 1994). Often, these introduced fungi and their pine hosts originated from different Northern Hemisphere continents, resulting in novel fungal-host pairings in their new ranges (Policelli et al., 2019; Hoeksema et al., 2020). Moreover, the introduction of soil inoculum selected for EcM fungal species that tolerated the transportation process and establishment in exotic plantation environments, inadvertently reduced fungal diversity in many introduced regions (Policelli et al., 2019; Hoeksema et al., 2020).

One notable case of such a co-introduction involves *Pinus radiata* and *Suillus luteus. Pinus radiata*, native to five isolated populations in coastal California and Mexico, is now the most extensively planted pine globally (Burdon et al., 2017). *Suillus luteus*, native to Eurasia where it is predominantly associated with *Pinus sylvestris*, is now found in pine plantations worldwide. *Suillus luteus* belongs to a suilloid clade of EcM fungi (also including the genus *Rhizopogon*) that specific to *Pinaceae* and has been closely associated with invasions of pines out of plantations throughout the Southern Hemisphere (Burdon and Chilvers, 1977; Richardson et al., 1994; Policelli et al., 2019). Introduced together to Australia as early as the 1860s, selective breeding of *P. radiata* began in the 1950s followed by widespread planting of improved seeds in the early 1970s (Wu et al., 2007). *Pinus radiata* now dominates the country’s softwood plantations and is grown in all states and territories except the Northern Territory (Wu et al., 2007). In Australia and several other countries, coinvasion of *P. radiata* and its fungal mutualists outside of plantations and into native habitats has caused concern from environmental and land management perspectives (Richardson et al., 1994; Policelli et al., 2019; Hoeksema et al., 2020).

Natural experiments like this can provide opportunities for valuable insights into phenomena such as rapid evolution, local adaptation, and coevolution (Rúa et al., 2016; Hoeksema et al., 2020). This study leverages this natural experiment to investigate how novel environments may have influenced the evolution of host compatibility in EcM fungal mutualists. Specifically, we used a cross-inoculation experiment to compare the compatibility of introduced Australian (AU) and native Czech (CZ) populations of *Suillus* luteus with native and introduced hosts, *Pinus sylvestris* and *Pinus radiata*. We aimed to a) investigate whether there is evidence of local adaptation in fungal isolates to their respective local host tree species, and b) determine how fungal origin affects the performance of those host tree species. We hypothesized that sympatric pairings would exhibit local adaptation, resulting in enhanced plant and fungal performance metrics. To address these objectives, we evaluated plant growth, survival, and morphology metrics, as well as fungal colonization intensity, providing proxies for assessing outcomes of evolutionary processes (Piculell et al., 2008).

## 2. Materials and Methods

### 2.1. Overview of Experimental Design

We conducted a cross-inoculation experiment using *Suillus luteus* collected from the introduced range (Australia (n=4) and New Zealand (n=1), n=5 isolates total) and the native range (Czech Republic (n=3) and Slovakia (n=2), n=5 isolates total) (supplementary table 1). These fungi were inoculated in all factorial combinations on genotypes of *Pinus radiata* selected for plantation forestry in Australia and New Zealand and on genotypes of *Pinus sylvestris* native to the Czech Republic. We also included non-inoculated controls, resulting in 22 different plant-fungus pairings. Each pairing was replicated four times, yielding a total of 88 experimental units. The entire experiment utilized soil collected from a site in New South Wales (NSW), Australia, where *P. radiata* and *S. luteus* are co-invading into native *Eucalyptus* forest from an adjacent pine plantation.

### 2.2. Seed, spore, and Soil Collection

*Pinus radiata* seeds were from two open-pollinated families sourced from New Zealand representing a typical genotype commonly used in Southern Hemisphere forestry (Sheffield’s Seeds, Lot 1824620, FM-6-c, FB-5-B12). *Pinus sylvestris* seeds were from an open-pollinated family collected from Krtiny in the Czech Republic (Sheffield’s Seeds-Lot 1830697, FB-2-H4), which is part of *S. lutues’* native range. *S. luteus* isolates were collected as spore prints from mushrooms found under *Pinus* species in Australia, New Zealand, the Czech Republic (Lobeš, Hradec Králové, and Bzenec-řívoz) and Slovakia (Šaštín-Stráže-Gazárka and Adamov) (supplementary table 1). Soil was collected underneath eight different mature *Eucalyptus racemosa* trees in Belanglo State Forest (NSW, Australia). Undecomposed litter was first removed, and soil was collected to a depth of approximately 10 cm.

### 2.3. Seed, Soil, and Spore Slurry Preparation and Inoculation

We surface sterilized seeds in a 10% bleach solution for two minutes, then rinsed them thoroughly in deionized water. After sterilization, seeds were soaked in water at 4ºC for 48 hours, drained, transferred to a sealed bottle with a moist paper towel, and cold stratified at 4ºC for 6 weeks.

Field soil was mixed with sand in a 1:1 ratio to improve drainage, then autoclaved (for one hour, at 121°C) twice with a 48-hour period in between, in plastic autoclave bags, in layers no more than 10cm thick. Because a lack of non-mycorrhizal soil microbes can substantially alter EcM fungal colonization of *Pinus* seedlings in experiments (Piculell et al. 2008), we added a microbial filtrate to the experimental soil. To create the filtrate, unsterilized Australian field soil was rinsed with approximately 2.7 L of water, which was then filtered consecutively through 1mm, 300µm, and 44µm filters to create a microbial wash. We then inoculated autoclaved Australian field soil with the microbial wash in batches with approximately 1.15 L of microbial wash per 7 L of soil.

For *S. luteus* inoculation treatments, we prepared spore slurries from spore prints scraped into deionized water, and the final spore concentration used in inoculation was 1.5×10^7^ spores per pot. Soil was inoculated in batches with 50ml of spore slurry and deionized water per 1.3L of soil.

### 2.4. Growth Conditions

After surface sterilization and cold stratification, pine seeds were planted approximately 1 cm below the soil surface in autoclave-sterilized potting soil. They were cultivated in a Conviron ATC40 growth chamber under 16-hour days (with 265 μmol/m2/s of light intensity at plant height) and 8-hour nights, maintaining a constant temperature of 22.0°C. After approximately three months, pine seedlings were transplanted into 3.81 × 20.955 cm (164 ml) cone-tainers filled with prepared Australian field soil, and then randomly distributed within their respective fungal inoculum groupings in the growth chamber. A 2.5 cm layer of sterilized sand was added on top of the field soil to serve as a barrier to prevent cross contamination of spores between pots. Throughout their growth stages, plants were watered 2-3 times per week with tap water.

### 2.5. Harvest and Data Collection

After 6 months, the plants were destructively harvested. Root systems were washed gently over a 2mm sieve to remove loose soil. Absolute EcM fungal colonization was assessed for each plant by counting colonized and uncolonized root tips. Dry weights of root and shoot material were recorded after 48 hours in a 60°C drying oven. Root length was estimated using the grid line intercept method (Tennant, 1975) 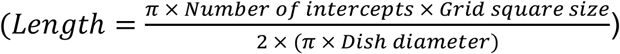 EcM fungal colonization intensity was calculated as the number of colonized root tips per unit of root length. Specific root length (SRL), a measure of root fineness, was calculated as estimated root length divided by root dry mass (cm/g). Root:shoot ratio was calculated as the ratio of root dry mass to shoot dry mass. EcM root tips were collected from a subset (n=6) of inoculated plants and were subjected to DNA extraction, direct PCR (with primers ITS1-F and ITS4), and Sanger sequencing (with conditions as described by Hoeksema et al., 2012), to verify the presence of *S. luteus*.

### 2.6. Data Analysis

All statistical analyses and plot building were conducted using RStudio (2024.04.1+748 “Chocolate Cosmos”) with the ggplot2, emmeans, dplyr, and tidyverse packages. Individual ANOVAs and *a priori* contrasts were performed to separately test whether each growth metric differed between fungal origins (Australia and Czech Republic) for each pine species (*P. radiata* and *P. sylvestris*), with the hypothesis that sympatric plant-fungal pairings would perform better than allopatric pairings. Fisher’s exact test was used to assess the effect of tree species and fungal origin on overall plant mortality.

## 3. Results

Very few uncolonized root tips were observed. Colonized EcM root tips on all inoculated plants exhibited typical suilloid morphology, and all root tips subjected to Sanger sequencing with fungal-specific primers revealed colonization by *S. luteus*. Mean total EcM colonization (# tips) and EcM colonization intensity (tips/cm) were both significantly higher when *P. sylvestris* was allopatrically paired with Australian (AU) fungal isolates than with sympatric Czech (CZ) isolates (p < 0.0001, p = 0.0003) (Figure 1a, 1b). *P. sylvestris* also had significantly higher shoot mass and total mass with AU fungi than with CZ fungi (p = 0.0005, p = 0.0014) (Figure 1e, 1d). In contrast, *P. radiata* exhibited higher root mass with AU fungi than with CZ fungi (p = 0.0108) (Figure 1f), while *P. sylvestris* showed a trend towards higher root:shoot ratio in AU fungal pairings compared to CZ (p = 0.0539) (Figure 1c).

**Figure 1.**
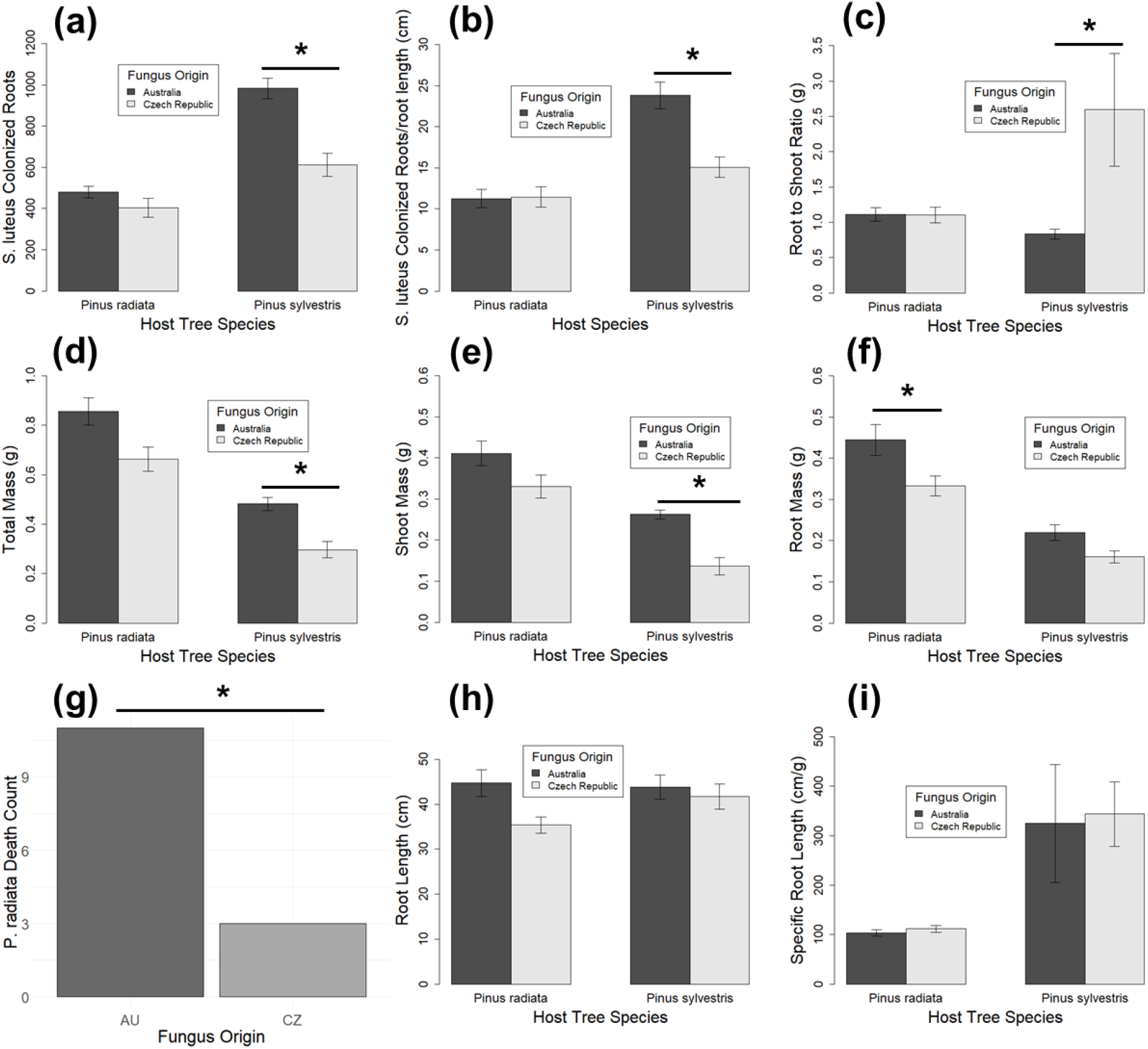
Metrics of plant and fungal performance: (a) *S. luteus* colonized root tips, (b) *S. luteus* colonization per root length, (c) Root:shoot ratio, (d) Total biomass, (e) Shoot biomass, (f) Root biomass, (g) *P. radiata* mortality count, (h) Root length, and (i) Specific root length. Asterisks indicate significant differences (p ≤ 0.05).

Specific root length (SRL) differed significantly between *P. radiata* and *P. sylvestris* (p < 0.0001), with *P. sylvestris* exhibiting higher SRL, or finer roots, compared to *P. radiata* (Figure 1i). *P. radiata* was the only species that experienced mortality (Fisher’s Exact Test, p = 0.00003078), and *P. radiata* mortality was significantly higher with AU fungi than with CZ fungi (Fisher’s Exact Test, p = 0.0187) (Figure 1g). With the exception of mortality and root mass, sympatric *P. radiata* pairings were not significantly different from allopatric pairings for any measure of plant or fungal performance. Non-inoculated treatments exhibited EcM colonization, likely due to movement of small numbers of *S. luteus* spores among pots, or due to colonization by EcM fungal spores included in the microbial wash, and thus were not included in analyses.

## 4. Discussion

We conducted a cross-inoculation experiment in which we tested how host species and fungal origin (native or introduced) affect compatibility of sympatric and allopatric pairs of pines and isolates of the introduced EcM fungus, *S. luteus*. Plant and fungal performance in the experiment was substantially affected by the pairings between plant and fungal genotypes, and the results largely support a hypothesis of local maladaptation in these interactions, with several metrics exhibiting stronger performance in allopatric pairings from both a plant and fungal perspective.

Despite an overall focus on local adaptation in the study of coevolved symbioses, local maladaptation has been increasingly recognized as an expected outcome in the coevolutionary process for many species interactions (Thompson et al., 2002). Maladaptation can arise from rapid environmental changes, the divergent evolutionary pressures of coevolution or competition (such as in arms races), or the introduction of maladaptive traits through gene flow. Moreover, research has shown that greater geographic overlap between partners diminishes the strength of EcM mutualism and increases the occurrence of local maladaptation (Nuismer et al., 2003; Karst et al., 2018). EcM fungi, often regarded as mutualists, operate along a mutualism-parasitism continuum influenced by environmental factors and the specific species involved in the interaction (Karst et al., 2008). Shifts along this continuum can lead to antagonistic evolution within coevolving mutualistic relationships resembling evolutionary arms races, where both the plant and fungus evolve to maximize their own performance and exploit the relationship. Consequently, these adaptations may diminish the mutualistic benefits provided by the fungus (Johnson et al., 1997). However, maladaptation does not necessarily signify the breakdown of a mutualistic relationship. Hosts may maintain low-quality partners in variable environments, even in the presence of alternatives, because they can provide short term benefits (Moeller and Neubert, 2016). For instance, *S. luteus* has been shown to facilitate pine invasion on its own, not because it is an ideal partner, but due to its long-range dispersal traits and a long-lived spore bank that extends beyond the existing range of the invading pines (Hayward et al., 2015). This reinforces the concept that coevolution is an ongoing process with periods of maladaptation and adaptation, rather than a static state of perfect mutualism.

Maladaptation in the EcM symbiosis between *S. luteus* and its native and exotic pine hosts is implied by several aspects of our results. First, *P. radiata* sympatrically paired with AU fungal isolates demonstrated significantly increased mortality compared to the allopatric pairing (Figure 1i), with little variation in other measures of performance compared to allopatric pairings with CZ fungal isolates (Figure 1a-h). In addition, sympatric pairings of *P. sylvestris* with CZ fungal isolates had significantly elevated root:shoot ratio and decreased *S. luteus* colonization and colonization/root length compared to the allopatric pairing (Figure 1c,g), suggested that the plant is allocating more resources to root growth relative to shoot growth, potentially to compensate for inadequate nutrient acquisition due to poor colonization by CZ fungal isolates. Meanwhile, the intensity of EcM colonization was highest in allopatric pairings of *Pinus sylvestris* with AU fungi, and that allopatric pairing exhibited higher aboveground and total biomass compared to the sympatric pairing of *P. sylvestris* with CZ fungi.

Our observation of reduced relative plant and fungal performance in sympatric pairings suggests that fungal isolates may have evolved traits that are less beneficial or even somewhat harmful to their local host plants. In addition, given the long history of *P. radiata* as a global forestry staple, having undergone at least three generations of artificial selection in Australia alone (Wu et al., 2007; Burdon et al., 2017), it is possible that selective breeding for traits like growth and pathogen resistance may have indirectly diminished compatibility of *P. radiata* with certain species of EcM fungi. Such indirect effects of selective breeding on plant traits that govern interactions with symbiotic microbes has been demonstrated in other crop plants (Porter and Sachs, 2020), such as legumes with rhizobia and arbuscular mycorrhizal fungi (Liu et al., 2020). In addition, it has been shown that multiple different genes in the *Pinus* genome influence symbiotic compatibility with different EcM fungi (Hoeksema et al., 2012; Piculell et al., 2019), implying that pines may flexibly evolve to have different compatibility with different EcM fungi, such that interactions with some fungi may be altered while others remain unchanged or differently altered.

In contrast to the maladapted sympatric pairings, *P. sylvestris* paired with AU fungal isolates in allopatric pairings showed enhanced performance, exhibiting better growth metrics across various parameters. This suggests that AU fungi are more effective mutualists for *P. sylvestris* than their native CZ fungi, aligning with other research that has found tree seedlings can benefit from novel interactions with EcM fungi (Karst et al., 2018). This enhanced performance could be influenced by several biotic and abiotic factors, including soil conditions, which play a crucial role in the strength and direction of mycorrhizal mutualism (Hoeksema et al., 2010). Using Australian field soil in all treatments may have advantaged AU fungi, enhancing their performance and colonization efficiency compared to CZ fungi, particularly if AU fungal isolates have adapted to unique Australian soil conditions. However, the similarity in performance of *P. radiata* when paired with both AU and CZ fungi suggests that other factors, such as genetic variation and evolutionary history, may also influence the observed patterns. Another potential factor is root morphology, which has been found to influence competition for colonization as well as symbiotic function (Voller et al., 2024). *P. sylvestris* has finer, more branched roots compared to *P. radiata* (Figure 1b), which may allow AU fungi to colonize more extensively due to the increased availability of fine root tips. This may explain why *P. sylvestris* paired with AU fungi exhibited higher colonization compared to the sympatric pairing of AU fungi and *P. radiata*. However, this observation does not account for why CZ fungal isolates did not also exhibit higher colonization on *P. sylvestris*, nor why sympatric pairings of *P. sylvestris* with CZ fungal isolates performed worse than allopatric pairings, suggesting again that maladaptation may have also played a role in these outcomes.

Despite the findings of significant variation in multiple plant and fungal responses to our experimental treatments, this study is constrained by several limitations. First, our analysis was performed on seedlings and under controlled lab conditions, which may have limited our ability to detect local adaptation in later stage fitness components, such as seed production. However, the role of *S. luteus* (Hayward et al., 2015) and other suilloid EcM fungi (Baar et al., 1999; Taylor and Bruns, 1999; Horton, 2002; Ashkannejhad and Horton, 2006) in pine establishment, early succession, and invasion mean that interactions with seedling pine hosts are likely to be an important stage of the partnership for the fitness of both members of the symbiosis. Second, we have an incomplete understanding of the introduction history of the Australian *S. luteus* used in our experiment, and it is possible that the native source population of our AU fungal isolates may differ in key traits compared to the CZ source population. However, genetic studies comparing native European populations of *S. luteus* with introduced Australian populations have not revealed significant genetic variation (see Chapter 1 of this dissertation and Ke et al. 2024 unpublished manuscript). Third, we were unable to include native Czech Republic soil alongside Australian soil treatments, which may have resulted in an incomplete understanding if fungi have adapted more significantly to soil conditions rather than (or in addition to) host identity (Johnson et al., 2010). Finally, we encountered EcM colonization in our non-inoculated control plants and observed small numbers of EcM root tips colonized by non-target fungal species within experimental treatments, which may have contributed to unwanted noise in our plant and/or fungal performance data.

Overall, these results underscore the potentially significant impact of rapid evolution in *S. luteus* within its introduced range, influencing compatibility in the EcM symbiosis with pine hosts. This study highlights the intricate dynamics of coevolution and emphasizes the necessity of incorporating both local adaptation and maladaptation into our understanding of species interactions. Moreover, it underscores the role of both biotic and abiotic factors in shaping these complex relationships.

## Supporting information

Supplementary Table 1. Fungal isolate origins

## 6. Supplementary Material

Supplementary Table 1. Fungal isolate origins

